# AmyCo: the Amyloidoses Collection

**DOI:** 10.1101/479279

**Authors:** Katerina C. Nastou, Georgia I. Nasi, Paraskevi L. Tsiolaki, Zoi I. Litou, Vassiliki A. Iconomidou

## Abstract

Amyloid fibrils are formed when soluble proteins misfold into highly ordered insoluble fibrillar aggregates and affect various organs and tissues. The deposition of amyloid fibrils is the main hallmark of a group of disorders, called amyloidoses. Curiously, fibril deposition has been also recorded as a complication in a number of other pathological conditions, including well-known neurodegenerative or endocrine diseases. To date, amyloidoses are roughly classified, owing to their tremendous heterogeneity. In this work, we introduce AmyCo, a freely available collection of amyloidoses and clinical disorders related to amyloid deposition. AmyCo classifies 74 diseases associated with amyloid deposition into two distinct categories, namely 1) amyloidosis and 2) clinical conditions associated with amyloidosis. Each database entry is annotated with the major protein component (causative protein), other components of amyloid deposits and affected tissues or organs. Database entries are also supplemented with appropriate detailed annotation and are referenced to ICD-10, MeSH, OMIM, PubMed, AmyPro and UniProtKB databases. To our knowledge, AmyCo is the largest repository containing information about amyloidoses and diseases related to amyloid deposition. The AmyCo web interface is available at http://bioinformatics.biol.uoa.gr/amyco.

**Katerina C. Nastou** is a Ph.D. student in Bioinformatics, at the Department of Biology of the National and Kapodistrian University of Athens. She is currently working on computational analysis of membrane and amyloidogenic proteins as part of her Ph.D. thesis. Her research focuses on the study of protein-protein interactions and the visualization and analysis of biological networks for these protein families, on the computational prediction of protein structure and function and the design and development of biological databases.

**Georgia I. Nasi** is a Ph.D. student in Biophysics, at the Department of Biology of the National and Kapodistrian University of Athens and a second year student in the Bioinformatics Master’s Program, at the same department. She is currently conducting her Master’s thesis on the computational analysis and visualization of the interaction network of amyloidoses and proteins associated with these disorders. Her research for her Ph.D. focuses on biophysical and computational analysis of amyloidogenic proteins and peptide-analogues associated with amyloidoses.

**Dr. Paraskevi L. Tsiolaki** is a Biologist with an MSc in Bioinformatics and a PhD in Molecular Biophysics. She is currently working as a postdoctoral fellow in Dr V. Iconomidou’s group, Assist. Prof. at the National and Kapodistrian University of Athens and her research interests focus on Molecular Biophysics and Structural Biology. Her current research efforts have been directed towards identifying the structural characteristics that underlie the self-assembly mechanisms,governing amyloidogenicity. More specifically she works on the structure and self-assembly of different amyloidogenic proteins or amyloidogenic peptide-analogues, implicated with amyloidoses, utilizing biophysical and biochemical techniques. She is also working on the computational and structural analysis of the anomalous type of protein-protein interactions in protein aggregation, with particular focus on the development of novel therapeutic intervention strategies.

**Dr.Zoi Litou** works as a Special Laboratory Teaching Staff in “Bioinformatics-Biophysics” at the Section of Cell Biology and Biophysics, Department of Biology, National &Kapodistrian University of Athens. She has a PhD in Bioinformatics. She is currently working on computational analysis of membrane proteins focusing on the automated recognition and classification of single-spanning membrane proteins, CWPs, GPCRs and Ion channels. Biological Network Analysis, Prediction algorithms, Algorithm Visualization techniques in Bioinformatics, High throughput sequencing analysis and visualization, Clustering Analysis, Knowledge discovery, management and representation, Data integration, Chemoinformatics, Pharmacogenomics, Text Mining in Bioinformatics, Personalized Medicine, Parallel programming.

**Dr. Vassiliki A. Iconomidou** is an Assistant Professor of Structural Biology/Molecular Biophysics and a group leader of Biophysics and Bioinformatics Lab at the Department of Biology of the National and Kapodistrian University of Athens. Her research interests include: 1) Structural and self-assembly studies of fibrous proteins, which form extracellular, proteinaceous structures of physiological importance like lepidopteran, dipteran and fish chorions and arthropod cuticle, 2) Structural and self-assembly studies of silkmoth chorion peptide-analogues as novel self-assembled polymers with amyloid properties, aiming at the construction of novel biomaterials with extraordinary physical properties, 3) Experimental studiesof the role of a great variety of amyloidogenic (‘aggregation-prone’) peptides, predicted by our AMYLPRED prediction algorithm, in several widespread and also rare pathological amyloidoses. She had been visiting European Molecular Biology Laboratory (EMBL Heidelberg) for more than ten years, conducting research on molecular self-assembly focusing especially on functional, protective and pathological amyloids and amyloidoses, and she was there when she published the first article on natural protective amyloids. She is the author of 41 publications and 6 book chapters which focus mostly on functional and pathological amyloid studies.

## Introduction

Amyloidoses are a group of disorders typically characterized by the extracellular and/or intracellular deposition of misfolded protein aggregates, known as amyloid fibrils [1]. The deposition is mainly attributed to the abnormal transition of a soluble protein into highly ordered insoluble amyloid fibrils, which disrupts the normal tissue architecture and results in organ failure [2, 3, 4, 5]. Since their discovery, amyloidoses became the center of attention for the scientific community and underwent various classifications, mostly based on clinical observations [6, 7, 8]. A current classification system categorizes amyloidoses as either localized or systemic [8, 9, 10, 11], whereas other systems cluster amyloidoses as primary or secondary [12, 13], hereditary or acquired [6, 14] and parenchymal or mesenchymal [15].

“Amyloidosis” has long been used as a general term to describe a family of heterogenous pathologies, caused by the abnormal protein misfolding and deposition [6]. Nevertheless, amyloid deposition is recorded as a clinical abnormality, occurring in a broad range of devastating, well-known or less common, disorders [16, 17]. Striking examples include the neurodegenerative Alzheimer disease [18, 19], with approximately 30 million people affected throughout the world and Diabetes Mellitus type 2 [20, 21], a metabolic disorder responsible for more than 1.5 million deaths annually [22]. Interestingly, amyloidosis was also traced as a rare complication of other clinical conditions with high prevalence in the human population, like certain types of cancer [23, 24, 25, 26, 27, 28, 29] and other severe diseases [30, 31, 32, 33].

To date, there are a few studies that systematically gather data for amyloidoses and store them in biological databases. Such examples are the Mutations in Hereditary Amyloidosis database [14], AL-Base [34], AlzGene [35], PDGene [36], PDBase [37] and AD&FTDMD [38]. However, it is evident, by looking up all previous systematic studies,that the majority of currently available repositories are mainly dedicated to well-studied diseases, while there is a lack of systematic approaches for the most rare and understudied disorders, associated with amyloid deposition. This fact stresses the importance of creating an up-to-date database that can assemble and categorize the heterogeneous group of diseases, originating from the deposition of amyloid fibrils.

Considering all these aspects, in this study we introduce AmyCo, a comprehensive, well-annotated and updated online collection, which contains data for 74 disorders associated with amyloid deposition. AmyCo entries were thoroughly gathered and supplemented with detailed annotation. Hyperlinks to ICD-10 [39], MeSH [40], OMIM [41], PubMed [42], AmyPro [43] and UniProtKB [44] databases were also included as additional disease details. To our knowledge, AmyCo is currently the most extensive database for the remarkably heterogeneous group of diseases, emerging from the deposition of amyloid fibrils.

## Methods

### AmyCo Data collection and Classification

AmyCo disease entries were collected through an extensive literature search (March 2018, PubMed [42]). A total of 249 studies, indexed in PubMed, were used (Please refer to Supplementary File 1 for more details). Data were additionally manipulated and diseases were classified into two broad categories, based on the following scheme:

- Amyloidosis, when amyloid deposition is the main disease cause (e.g. AL amyloidosis, Alzheimer disease)
- Clinical conditions associated with amyloidosis, when amyloid deposition is detected but is neither the main nor the common disease cause (e.g. amyloid deposition in Waldenström’s macroglobulinemia)

### AmyCo Disease Nomenclature

Disease entry names were thoroughly reviewed and selected. The following nomenclature scheme was used for the main disease name:

- a MeSH name is assigned as a disease name when a disorder is recorded as a MeSH entry
- an ICD-10 name, is assigned as a disease name when there is no available MeSH entry, and
- the most common name of the disease found in the scientific literature is assigned as a disease name when a disease has neither a MeSH nor an ICD-10 entry.

Common or less common disease names were also included as alternative disease terms. The International Society of Amyloidosis (ISA) nomenclature was additionally added for disease entries related to the majority of proteins that have been recorded as amyloid fibril proteins in human according to the ISA [45].

### Additional AmyCo features

Each database entry was manually annotated with major protein components (causative protein) and minor protein components. The latter were especially included in the “Amyloidosis” category in an attempt to correlate distinct protein components that happen to be co-deposited in the amyloid plaques of one or more conditions [46, 47]. Co-deposited components are experimentally verified by either immunohistochemistry, MS-based proteomics, staining or imaging techniques [46, 47, 48, 49, 50]. Protein nomenclature, used throughout the database, follows the modern guidelines suggested by the ISA [45]. In addition to this feature, diseases belonging to the “Clinical conditions associated with amyloidosis” category were annotated with a related “Amyloidosis” category, utilizing a hyperlinked section. This AmyCo feature unravels the relationship between pathologies at a disease level, based on existing literature data.

For each entry, information about the respective disease and associated proteins is provided, along with literature references (PubMed [42]), cross-references to major publicly available disease databases -when available-(MeSH [40], ICD [39], OMIM [41]) and protein databases (UniProt [44], AmyPro [43]). Tissues or organs affected by amyloid deposits were also collected from the literature and added as additional disease details [42]. Entries, which have MeSH records, were further enriched with a MeSH description [40]. A supplementary disease subdivision was embed, based on the well-established ICD-10 classification scheme [39].

### AmyCo Implementation

A web application for AmyCo has been created with a two-layer approach; a mySQL database system and a Node.js application server. The first layer consists of a mySQL database management system, with all disease and protein data stored in a relational database. The second layer is a Node.js application server that receives user queries to the database and returns data to the web browser. The web interface is based on modern technologies (HTML5, CSS3 and Javascript) and can be viewed from any screen size (desktop, tablet or mobile). A CytoscapeJS [51] viewer is integrated for the visualization of the association between diseases and protein components of amyloid deposits.

## Results & Discussion

The deposition of amyloid fibrils has been linked to the development of a broad class of life-threatening diseases, called amyloidoses [1]. While amyloid fibril deposition has been recorded as a major and/or minor complication in a number of pathological conditions, it still remains unclear whether it is the cause or the consequence of these diseases [16]. Despite intensive studies during the past decades in the field of amyloid research, until now there is a lack of a standard classification for amyloidoses. In this work, we created AmyCo, a novel manually curated collection of amyloidoses and clinical conditions associated with amyloid deposition. AmyCo displays a novel classification system for these rare disorders and provides links to important literature references and other significant disease databases. To the best of our knowledge, AmyCo collects the largest number of disorders (74) related to amyloid deposition and associates proteins acting as principal causative disease agents (83). A detailed comparison between AmyCo and other available resources is presented in Table 1.

**Table 1.**
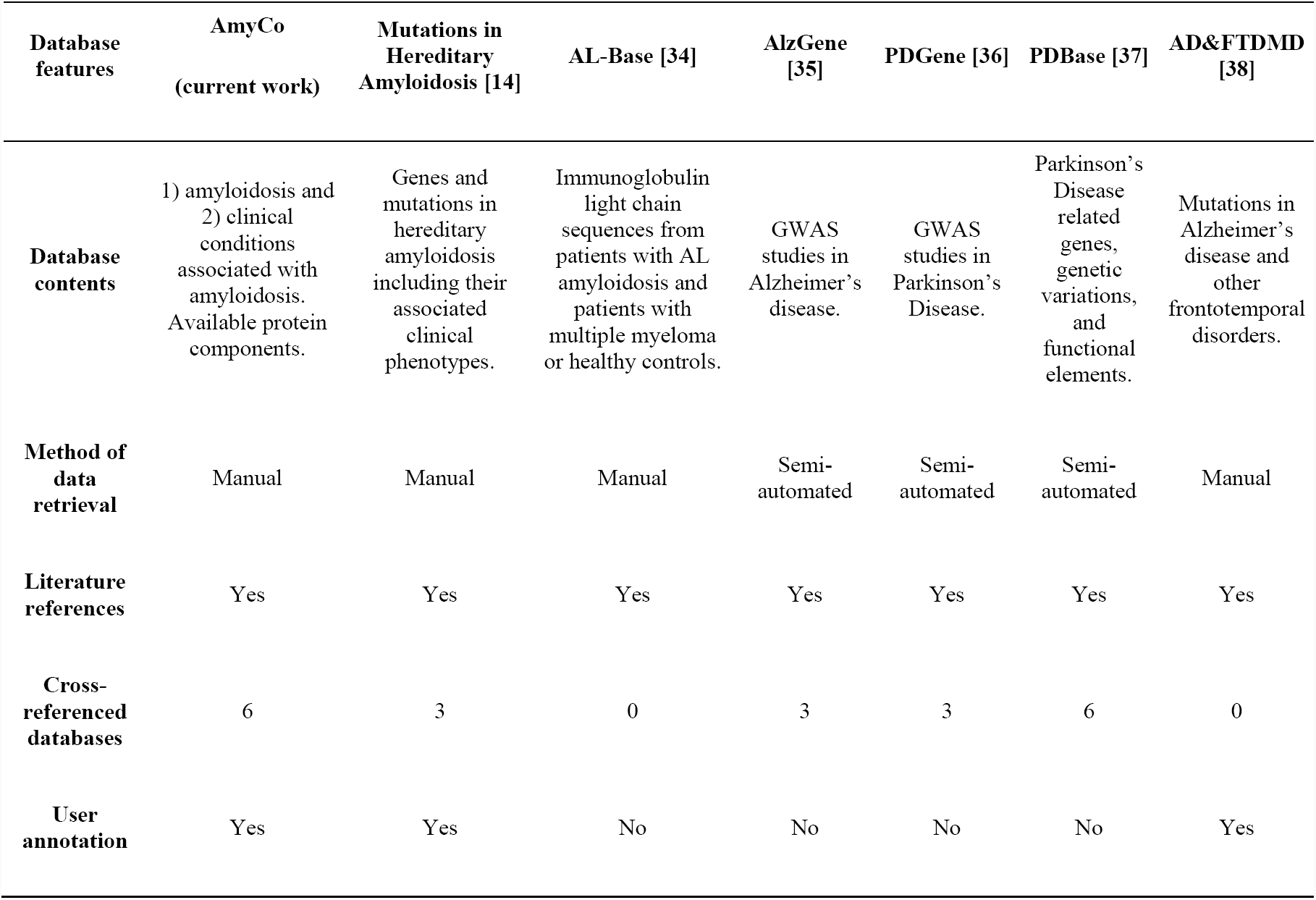
A comparison between AmyCo and other databases that include information of diseases, associated with amyloid deposition.

One of the main challenges during the creation of AmyCo was to manually filter the plethora of available information, form the final non-redundant dataset of diseases and clarify nomenclatures for both the diseases and their associated proteins. In a similar way, an interesting approach to group amyloidosis, based on the type of precursor proteins that form insoluble amyloid fibrils, was introduced by Misumi and Ando, back in 2014 [52]. In turn, our final classification scheme divided 74 diseases into two distinct categories, namely “amyloidosis” and “clinical conditions associated with amyloidosis” (Please see Materials and Methods for more details). This classification scheme allowed us to unify current and older nomenclatures, accurately allocate alternative disease names, and subsequently, gather relevant knowledge from both the scientific literature [42] and other more generic databases [39, 40, 41]. Literature data reveal that when it comes to amyloidoses there is a variable use of terms [8, 53], despite the efforts of the ISA nomenclature committee to establish a common classification system [6, 45, 54]. Alternative disease terms were included in an attempt to ensure the consistency of our database and improve the overall information flow. Universal MeSH and ICD-10 terms were assigned as the main disease name, following the scheme described in the Materials and Methods section. Thus, AmyCo is a valuable reference for anyone using contemporary nomenclature or older disease designations that are still extensively in use in the literature.

At the same time, 83 experimentally validated protein components of amyloid deposits were assigned to each of the diseases (Figure 1). The ISA classification system was used as a reference for the nomenclature of these protein components. The co-existence of proteins in amyloid deposits gained a lot of attention due to its established connection with protein aggregation [17, 55]. A variety of experimental techniques, direct or indirect, are used to capture this common phenomenon [46, 47, 48, 49, 50]. Such examples are the Gerstmann-Straussler-Scheinker Syndrome, where amyloid PrP (APrP) and Aβ are both found in amyloid plaques [49], or the Alzheimer’s disease, in which many proteinaceous components have been recorded as co-deposits of the amyloid plaques [48, 56, 57]. Co-deposition of proteins, occurred either through cross-seeding [17, 55] or cross-inhibition [17, 55] is an extremely important detail for these rare diseases, as it could be a key starting point towards associating the majority of amyloid-related diseases and understanding the inevitable cascade of protein aggregation.

**Figure 1.**
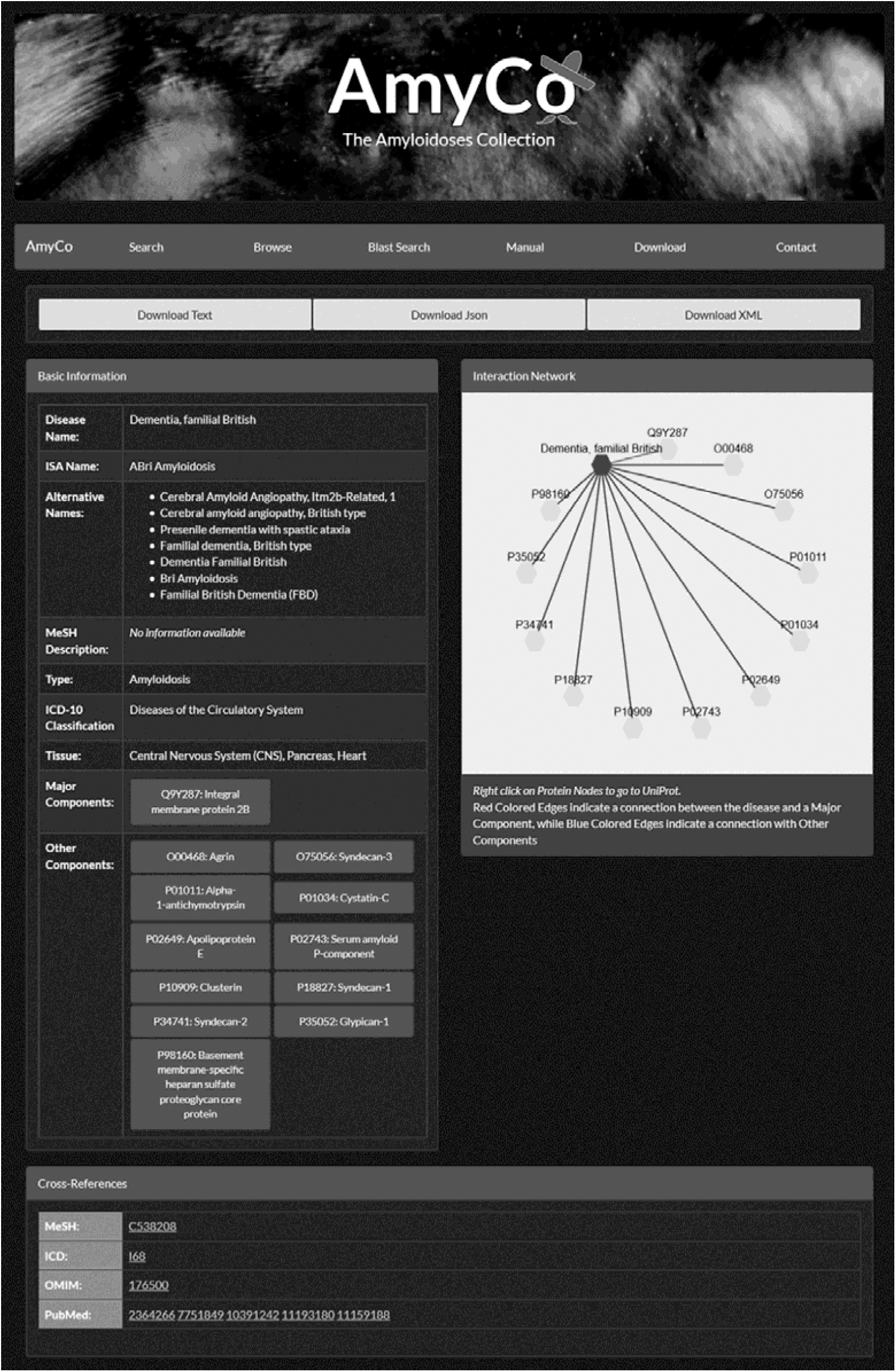
An AmyCo disease entry. The user can access basic information about the disease, including the disease name, a MeSH short description, an ICD-10 classification and the major or other protein component, related to the disease. Cross-references to other databases are also provided. All data are available for download in text, JSON and XML formats.

Apart from the systematic data collection from the scientific literature, which will be performed in a regular basis in order to maintain AmyCo records, an interesting option is the feature that allows the scientific community contribution. User contribution is an inseparable component of biological databases that wish to stay updated and incorporate the ever-growing biological knowledge [58, 59]. Furthermore, community annotation allows the synergy between curators and researchers, helps towards the maintenance of a repository and promotes interdisciplinary collaborations [60]. In the case of AmyCo, user annotation is a feature of utmost importance, since this collection was based upon the entire scientific literature (Supplementary File 1) and thus, some data may be falsely filtered out or certain biases may be reflected during the disease collection. More importantly, there is an excessive amount of information about amyloidoses that is generated daily by the community and added to the amyloid research vault. A submission form, provided in the contact page of the AmyCo v1.1 web application, allows user annotation and welcomes the insertion of comments or new data. This utility renders AmyCo a valuable tool for the interaction of the scientific community dealing with amyloidoses.

### User interface and website features

The AmyCo database has a user-friendly interface that offers convenient ways to gain access into its data. The ‘Home’ page provides a short description and database statistics. From the navigation bar, at the top of every page, users can either perform searches or browse the database contents. A ‘Manual’ page explaining the functionalities of AmyCo, a ‘Contact’ page with author contact information and a submission form for user contribution are also available.

Search queries utilize either simple protein and disease names or the more complex UniProt ACs/IDs [44] and HUGO Gene Names [61]. Terms can be used indiscriminately by the user as active search keywords, provided that are spelt correctly (Please refer to Supplementary File 2 for more details). Results can be sorted by using our disease classification system. While browsing AmyCo, a user can have access to all disease entries or filter them by disease category and/or ICD-10 classification [39]. Results retrieved from both browsing and searching the database are displayed in tables. Direct links to disease entry pages are given at the end of each row. An additional BLAST search tool[62] is integrated for running protein BLAST searches against the database, using as input one or more FASTA formatted sequences [63].

AmyCo is currently available in three formats (text, XML and JSON) and is supported for download by the ‘Download’ button, at the top navigation bar. The AmyCo manual is also provided as a separate supplementary file (Supplementary File 2).

### Disease Entries

Database entries are generated dynamically via browsing, searching or through direct URL links. As shown in Figure 1, direct links for data downloading are provided on the top of each page. A table displaying all entry sections (e.g. disease name, ISA name, alternative name etc.) is also available. CytoscapeJS [51] is used to visualize all the relationships between a disease and its associated proteins in a comprehensive functional context [64]. Detailed information about each related protein can be also viewed by pressing the respective button. Major protein components are also supplemented with AmyPro links [43] and useful literature references, whereas experimental evidence is also provided for all the co-deposited amyloid components. External interconnections with literature references, disease repositories and protein databases enhance each disease entry with significant details (Please refer to the Supplementary File 2).

## Conclusions

AmyCo assembles and categorizes the heterogeneous group of diseases, associated with the deposition of amyloid fibrils. Our novel database provides a uniform access to data recorded in different literature sources and classifies 74 diseases into two distinct categories, namely 1) amyloidosis and 2) clinical conditions associated with amyloidosis. The added value of detailed literature references and the manual annotation feature render AmyCo a unique database for diseases associated with amyloid deposition. It is hoped that this approach will aid both clinical scientists and researchers, in the need of a comprehensive resource, referencing biological information on amyloidoses. AmyCo is available at http://bioinformatics.biol.uoa.gr/amyco

## Supporting information

## Acknowledgements

The authors thank the National and Kapodistrian University of Athens for support. The authors would also like to thank the anonymous reviewers and the handling editor for their valuable comments and constructive criticism.

## Author Contributions

Study design: KCN, GIN, PLT, ZIL, VAI; Conceptualization: ZIL, KCN, VAI; Literature and Database search: GIN; Database design and development: KCN; Data Curation: PLT, GIN, KCN, ZIL; Web Application Design: KCN; Web Application Quality Assurance: PLT, GIN, KCN, ZIL, VAI; Writing - original draft: KCN, GIN, PLT; Writing - review and editing: GIN, PLT, KCN, ZIL, VAI; Supervision: VAI; Funding Acquisition: VAI.

