## Supplementary material for "AmyCo: the Amyloidoses Collection"

*Supplementary File 1*

### AmyCo: the Amyloidoses Collection

Katerina C. Nastou[#], Georgia I. Nasi[#], Paraskevi L. Tsiolaki[#], Zoi I. Litou and Vassiliki A. Iconomidou*

*Section of Cell Biology and Biophysics, Department of Biology, School of Sciences, National and Kapodistrian University of Athens, Panepistimiopolis, Athens 157 01, Greece*

[#]These authors contributed equally.

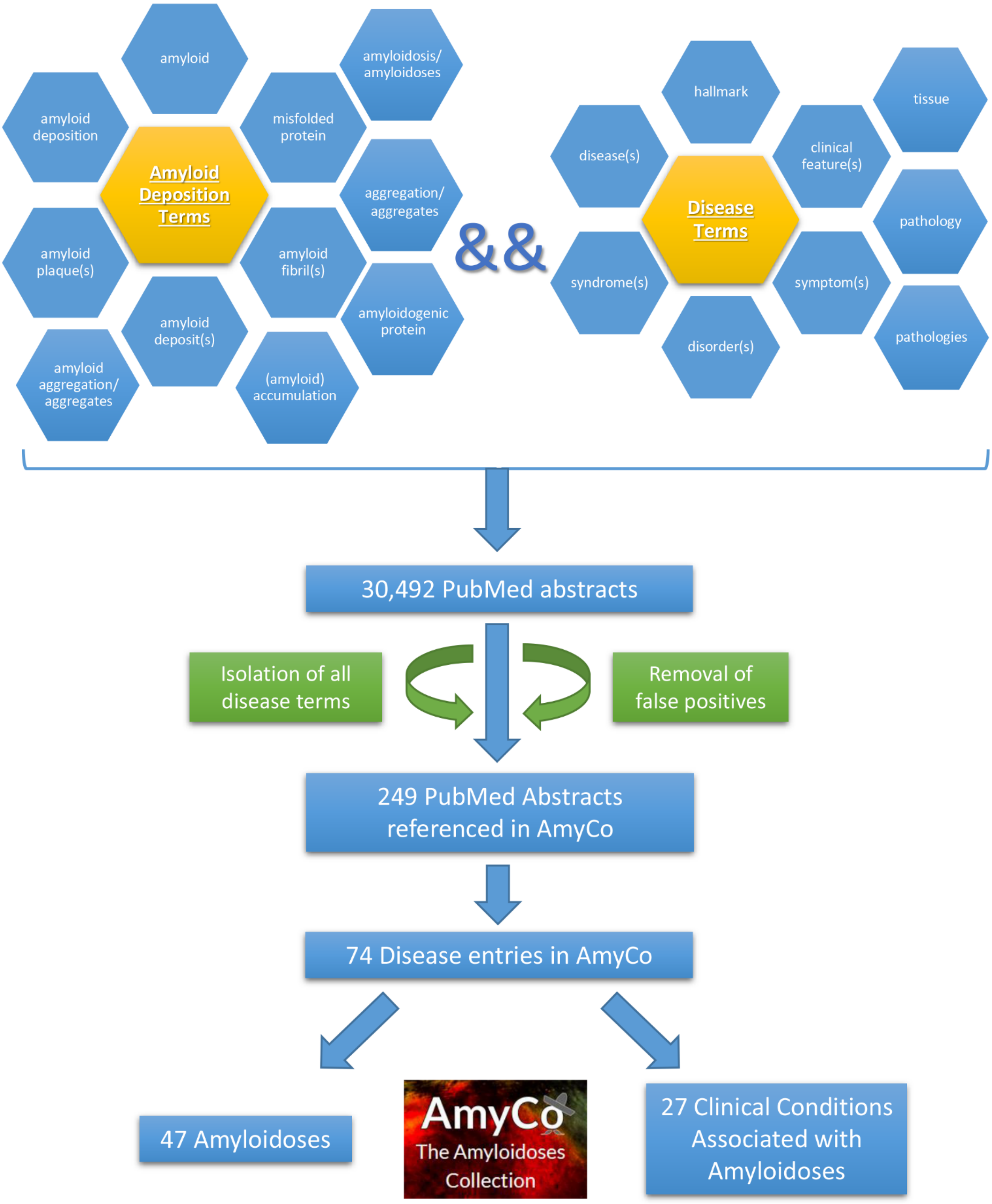


**Supplementary Figure 1.** ***AmyCo data collection***. An initial search in PubMed was carried out for articles, combining terms related to "amyloid deposition" and "disease". This search returned 30,492 PubMed Abstracts. Due to the large number of the false positive hits, results were manually filtered; abstracts containing disease terms were gathered, whereas abstracts having false positive terms were removed. This filtering and manipulation led to 249 PubMed Abstracts referring to amyloidoses and diseases associated with amyloid deposition used for the creation of AmyCo. AmyCo (http://bioinformatics.biol.uoa.gr/amyco) classifies 74 diseases associated with amyloid deposition into two categories, namely 1) amyloidoses and 2) clinical conditions associated with amyloidoses.
