## Supplementary material for "AmyCo: the Amyloidoses Collection"

*Supplementary File 2*

### AmyCo: the Amyloidoses Collection

Katerina C. Nastou[#], Georgia I. Nasi[#], Paraskevi L. Tsiolaki[#], Zoi I. Litou and Vassiliki A. Iconomidou*

*Section of Cell Biology and Biophysics, Department of Biology, School of Sciences, National and Kapodistrian University of Athens, Panepistimiopolis, Athens 157 01, Greece*

[#]These authors contributed equally.

### AmyCo

The Amyloidoses Collection

**Welcome to *AmyCo*, a freely available, literature-curated collection of amyloidoses and other pathological conditions associated with the deposition of amyloid fibrils.**

Manual Contents


1. Home
2. Search
3. Browse
4. Disease Entry
5. BLAST Search
6. Download
7. Contact
8. Database Technologies

#### Home

In order to visit *AmyCo*, the user should enter the following address: [http://bioinformatics.biol.uoa.gr/AmyCo/](http://bioinformatics.biol.uoa.gr/AmyCo/). The page contains a short description of the repository and the *AmyCo* statistics.

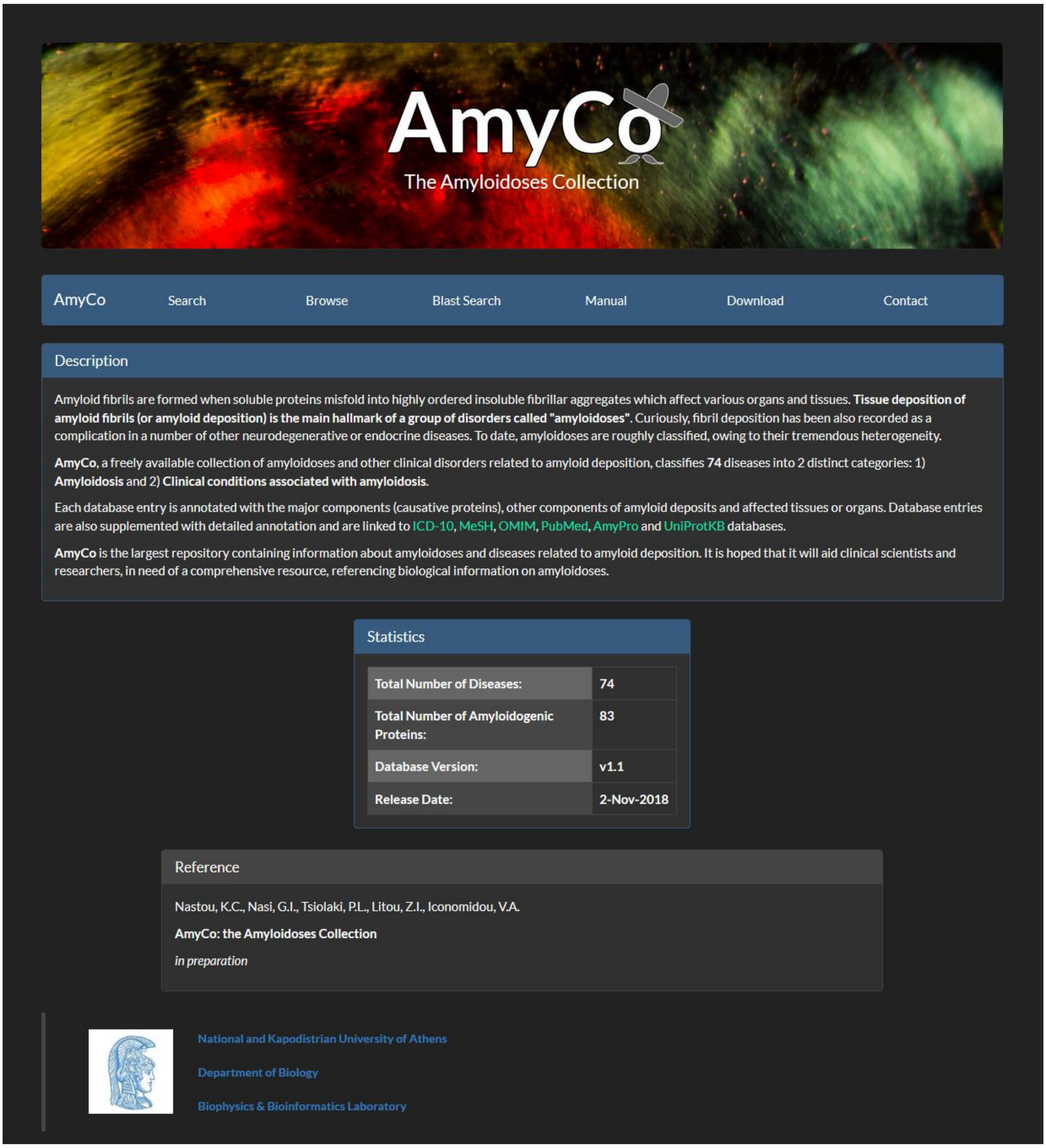

#### Search

The search tab allows the navigation of the database. A form with multiple options appears.

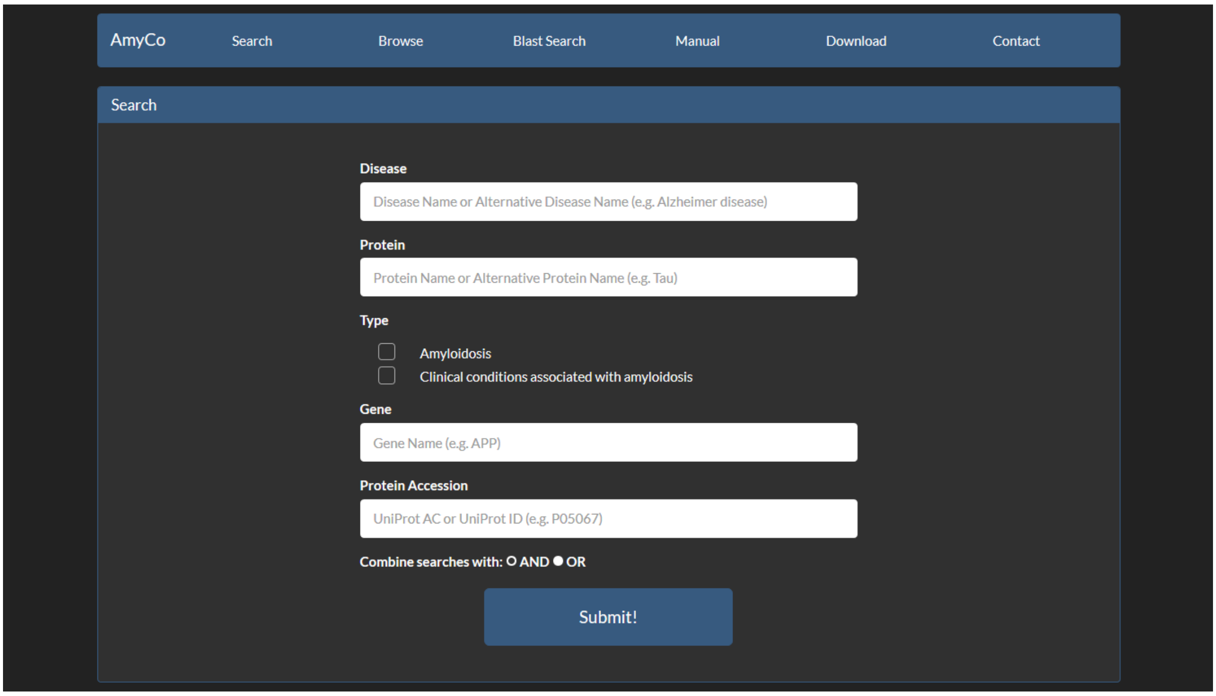


The search options are:

- by Disease Name

  The user may use a Name or an Alternative Disease Name, including the ISA name when available (e.g. Alzheimer Disease or Alzheimer Syndrome)

- by a Protein Name associated with a disease

  Proteins can be either a major or a minor (other) component.

- by Disease Type

  AmyCo Disease entries are classified into two categories: **1) Amyloidosis**, when amyloid deposition is the main disease cause (e.g. AL amyloidosis) or amyloid deposits are present in tissue and organs (e.g. Alzheimer disease), and **2) Clinical conditions associated with amyloidosis**, when amyloidosis is a clinical feature of a disease or a syndrome (e.g. amyloid deposition in Waldenström's macroglobulinemia)

- by Gene Name (e.g. APP)

- by Protein Components based on a UniProt Accession or a UniProt Identifier (e.g. P05067)

AmyCo
The Amyloidoses Collection

The search, based on disease name, protein name and gene name does not require specific words. For example a user enters the word "apo" in the **disease search field**.

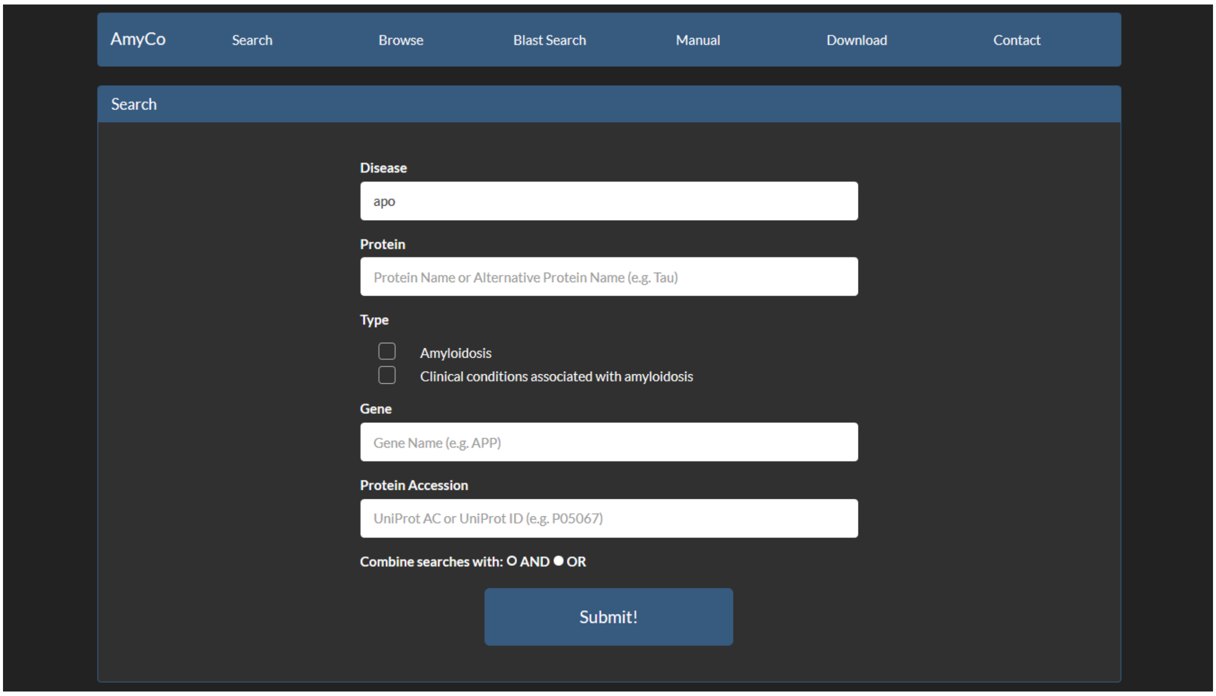


The result is all diseases containing the word "apo" in the **disease search field**.

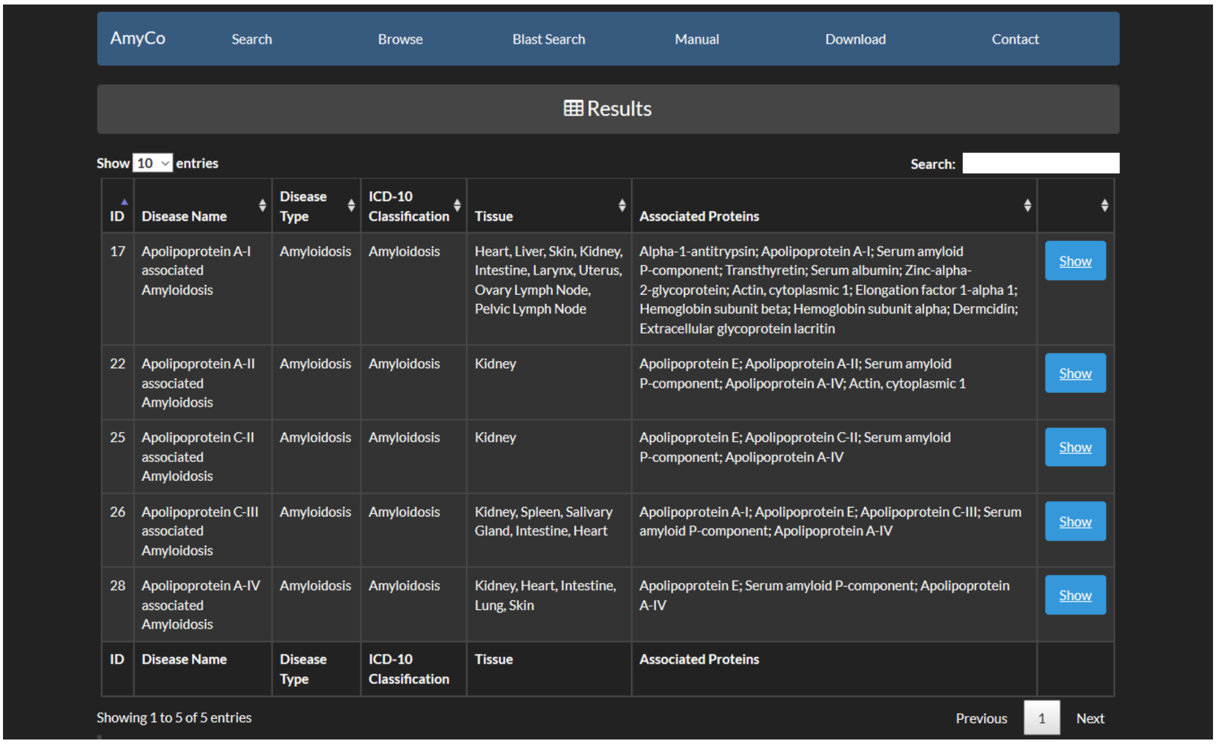


| ID | Disease Name | Disease Type | ICD-10 Classification | Tissue | Associated Proteins | |
|---|---|---|---|---|---|---|
| 17 | Apolipoprotein A-I associated Amyloidosis | Amyloidosis | Amyloidosis | Heart, Liver, Skin, Kidney, Intestine, Larynx, Uterus, Ovary Lymph Node, Pelvic Lymph Node | Alpha-1-antitrypsin; Apolipoprotein A-I; Serum amyloid P-component; Transthyretin; Serum albumin; Zinc-alpha-2-glycoprotein; Actin, cytoplasmic 1; Elongation factor 1-alpha 1; Hemoglobin subunit beta; Hemoglobin subunit alpha; Dermcidin; Extracellular glycoprotein lacritin | Show |
| 22 | Apolipoprotein A-II associated Amyloidosis | Amyloidosis | Amyloidosis | Kidney | Apolipoprotein E; Apolipoprotein A-II; Serum amyloid P-component; Apolipoprotein A-IV; Actin, cytoplasmic 1 | Show |
| 25 | Apolipoprotein C-II associated Amyloidosis | Amyloidosis | Amyloidosis | Kidney | Apolipoprotein E; Apolipoprotein C-II; Serum amyloid P-component; Apolipoprotein A-IV | Show |
| 26 | Apolipoprotein C-III associated Amyloidosis | Amyloidosis | Amyloidosis | Kidney, Spleen, Salivary Gland, Intestine, Heart | Apolipoprotein A-I; Apolipoprotein E; Apolipoprotein C-III; Serum amyloid P-component; Apolipoprotein A-IV | Show |
| 28 | Apolipoprotein A-IV associated Amyloidosis | Amyloidosis | Amyloidosis | Kidney, Heart, Intestine, Lung, Skin | Apolipoprotein E; Serum amyloid P-component; Apolipoprotein A-IV | Show |
| ID | Disease Name | Disease Type | ICD-10 Classification | Tissue | Associated Proteins | |

All searches can be combined with logical operators (AND/OR), in order to make the search result as specific as possible.

For example if we proceed to the following combined search:

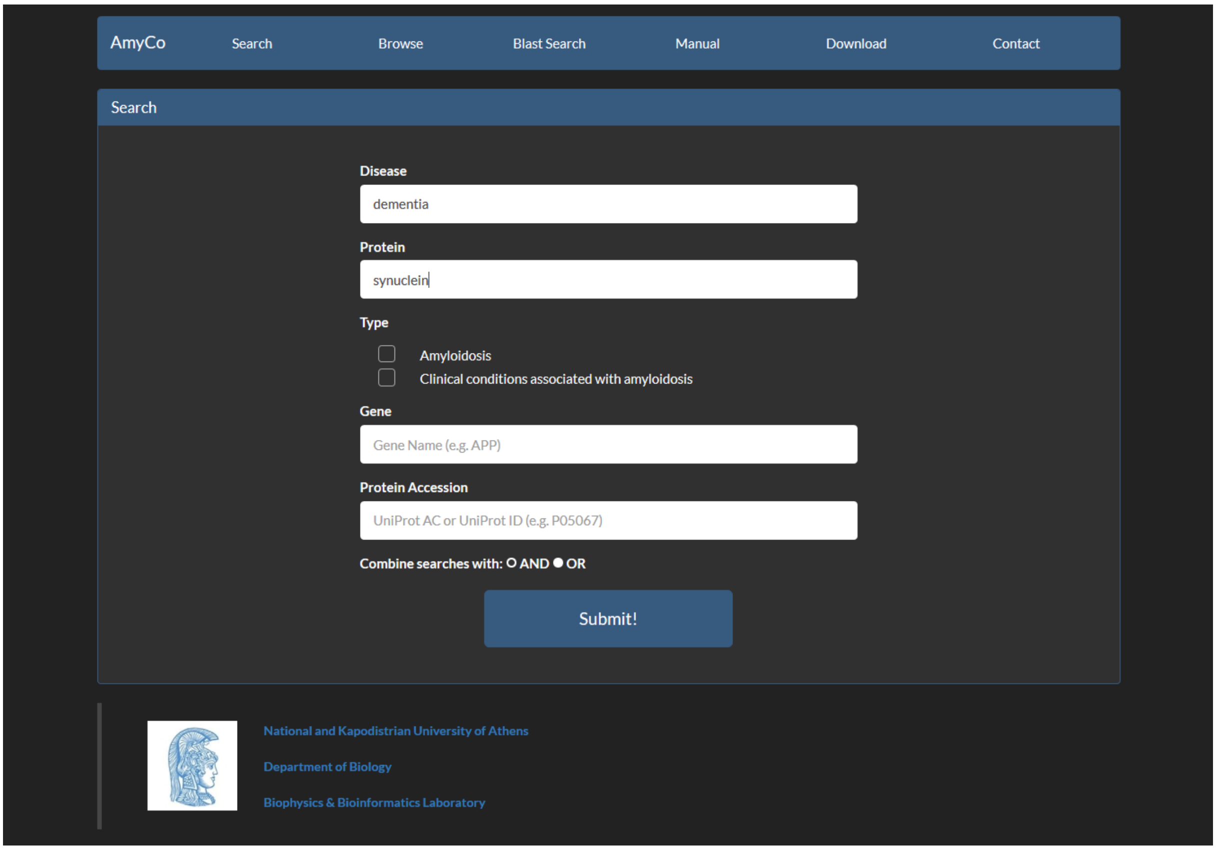


We will get only the disease, which is classified as dementia and is associated with synuclein.

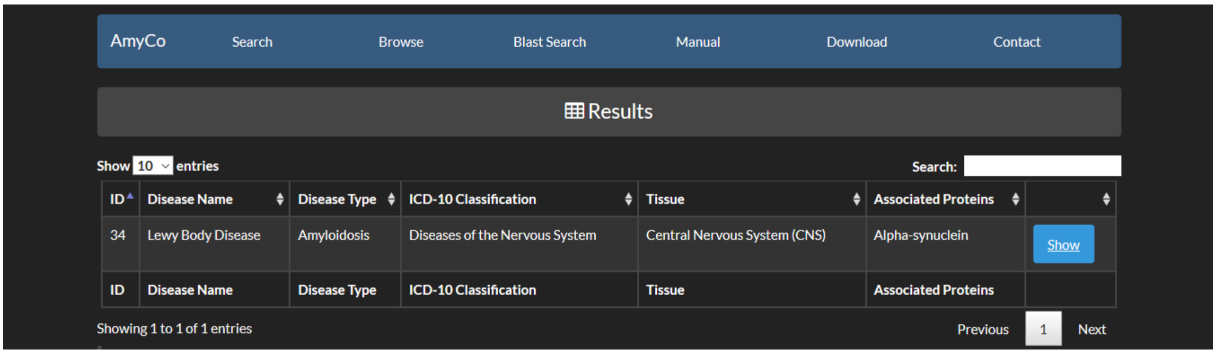

#### Browse

The **browse tab** allows the browsing of the database. User can apply filters and browse the database by selecting the disease Type or/and the ICD-10 Classification.

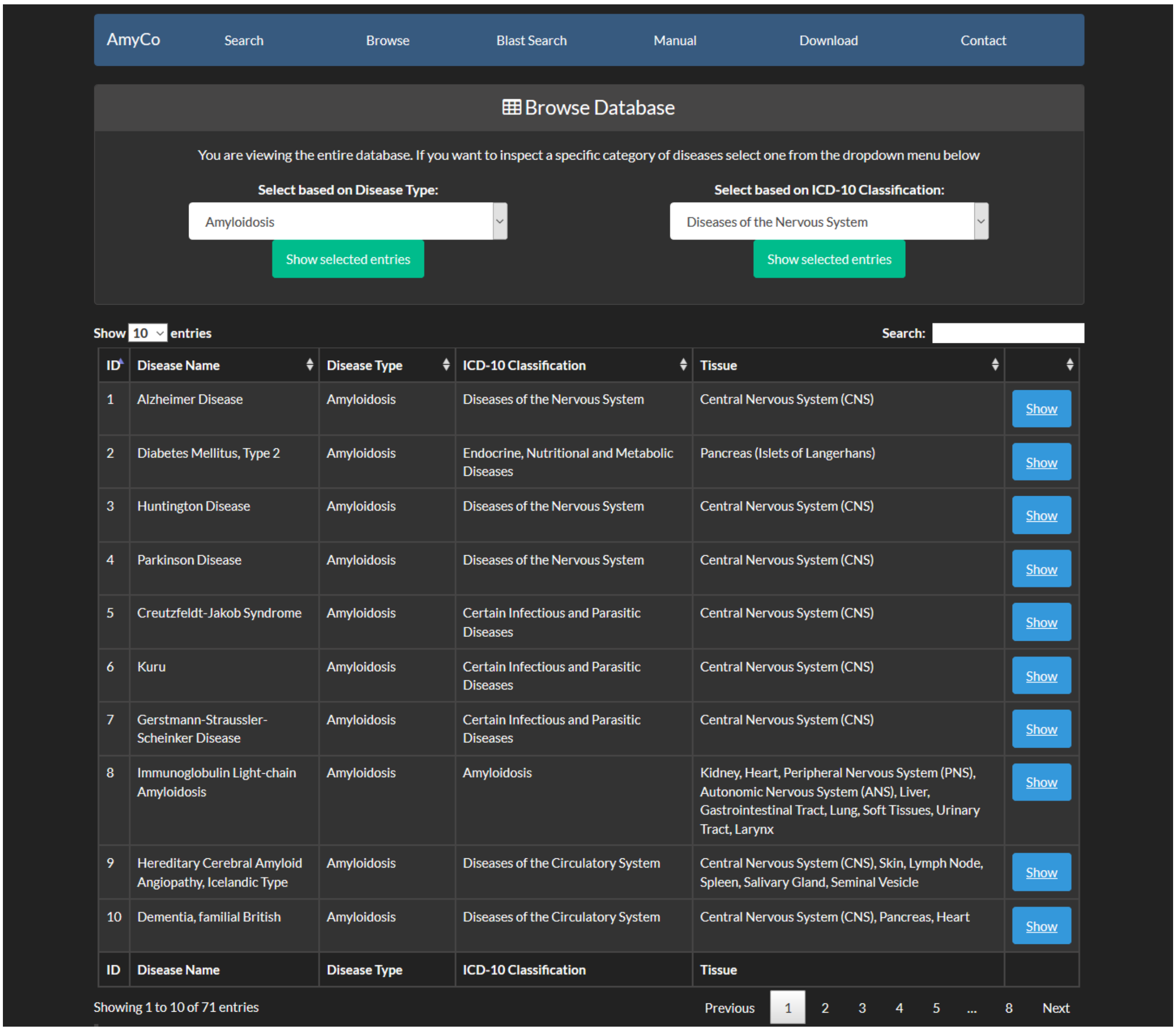
AmyCo Search Browse Blast Search Manual Download Contact

Browse Database

You are viewing the entire database. If you want to inspect a specific category of diseases select one from the dropdown menu below

Select based on Disease Type: Amyloidosis — Show selected entries

Select based on ICD-10 Classification: Diseases of the Nervous System — Show selected entries

Show 10 entries — Search:

| ID | Disease Name | Disease Type | ICD-10 Classification | Tissue | |
|---|---|---|---|---|---|
| 1 | Alzheimer Disease | Amyloidosis | Diseases of the Nervous System | Central Nervous System (CNS) | Show |
| 2 | Diabetes Mellitus, Type 2 | Amyloidosis | Endocrine, Nutritional and Metabolic Diseases | Pancreas (Islets of Langerhans) | Show |
| 3 | Huntington Disease | Amyloidosis | Diseases of the Nervous System | Central Nervous System (CNS) | Show |
| 4 | Parkinson Disease | Amyloidosis | Diseases of the Nervous System | Central Nervous System (CNS) | Show |
| 5 | Creutzfeldt-Jakob Syndrome | Amyloidosis | Certain Infectious and Parasitic Diseases | Central Nervous System (CNS) | Show |
| 6 | Kuru | Amyloidosis | Certain Infectious and Parasitic Diseases | Central Nervous System (CNS) | Show |
| 7 | Gerstmann-Straussler-Scheinker Disease | Amyloidosis | Certain Infectious and Parasitic Diseases | Central Nervous System (CNS) | Show |
| 8 | Immunoglobulin Light-chain Amyloidosis | Amyloidosis | Amyloidosis | Kidney, Heart, Peripheral Nervous System (PNS), Autonomic Nervous System (ANS), Liver, Gastrointestinal Tract, Lung, Soft Tissues, Urinary Tract, Larynx | Show |
| 9 | Hereditary Cerebral Amyloid Angiopathy, Icelandic Type | Amyloidosis | Diseases of the Circulatory System | Central Nervous System (CNS), Skin, Lymph Node, Spleen, Salivary Gland, Seminal Vesicle | Show |
| 10 | Dementia, familial British | Amyloidosis | Diseases of the Circulatory System | Central Nervous System (CNS), Pancreas, Heart | Show |
| ID | Disease Name | Disease Type | ICD-10 Classification | Tissue | |

Showing 1 to 10 of 71 entries — Previous 1 2 3 4 5 ... 8 Next

At first all entries appear. Each page shows 10 entries by default but this can be altered from the dropdown menu at the top left corner of the table to 25, 50 or all entries. Moreover, users can perform non-specific searches using the search option at the top right corner of the data table.

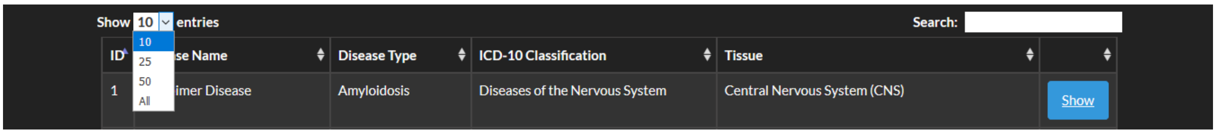

Filters and browsing can be applied according to the classification below.

**Disease type and ICD-10 classification**

| | Disease Type | ICD-10 Classification |
|---|---|---|
| 1 | **Amyloidosis** | Amyloidosis |
| | | Endocrine, Nutritional and Metabolic Diseases |
| | | Diseases of the Circulatory System |
| | | Chromosomal Abnormalities |
| | | Diseases of the Nervous System |
| | | Neoplasms |
| | | Certain Infectious and Parasitic Diseases |
| | | Diseases of the Eye and Adnexa |
| 2 | **Clinical conditions associated with amyloidosis** | Endocrine, Nutritional and Metabolic Diseases |
| | | Diseases of the Respiratory System |
| | | Diseases of the Musculoskeletal System and Connective Tissue |
| | | Neoplasms |
| | | Diseases of the Skin and Subcutaneous Tissue |
| | | Diseases of the Digestive System |
| | | Diseases of the blood and blood-forming organs and certain disorders involving the immune mechanism |
| | | Other |

For example, if the user selects **Amyloidosis** as the disease type and **Neoplasms** as the ICD-10 classification and presses *Show Selected Entries*.

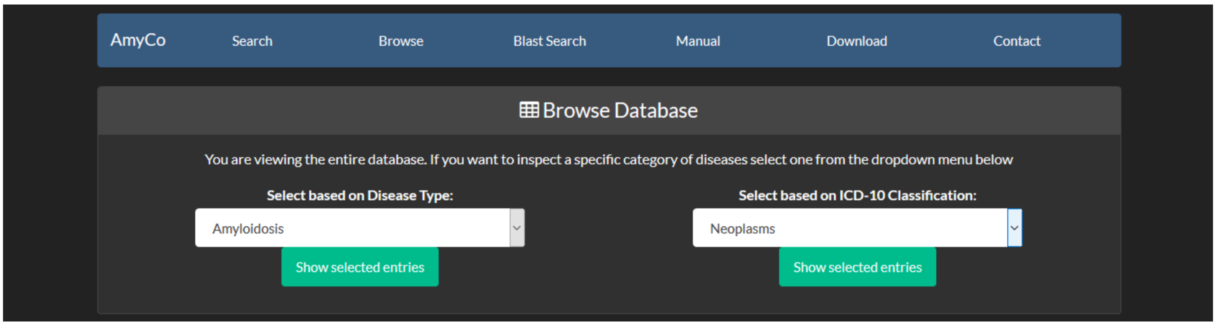


The selected diseases are:

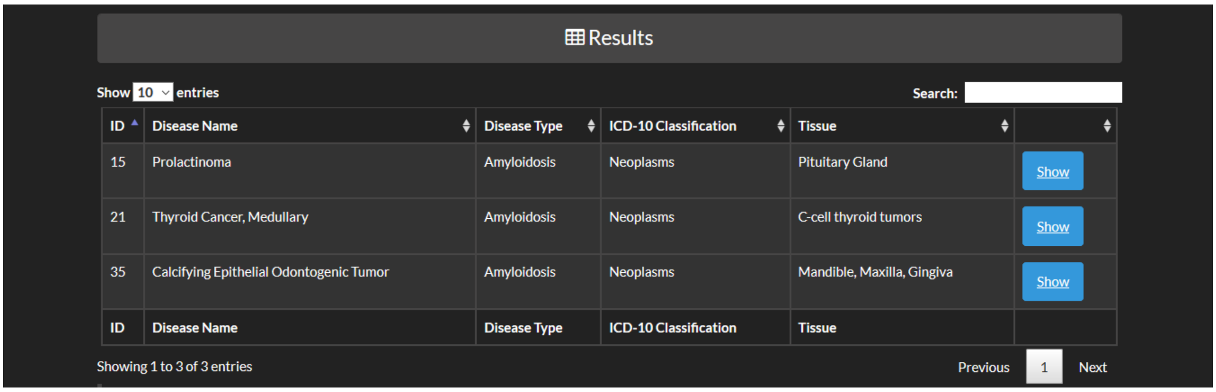


| ID | Disease Name | Disease Type | ICD-10 Classification | Tissue | |
|---|---|---|---|---|---|
| 15 | Prolactinoma | Amyloidosis | Neoplasms | Pituitary Gland | Show |
| 21 | Thyroid Cancer, Medullary | Amyloidosis | Neoplasms | C-cell thyroid tumors | Show |
| 35 | Calcifying Epithelial Odontogenic Tumor | Amyloidosis | Neoplasms | Mandible, Maxilla, Gingiva | Show |
| ID | Disease Name | Disease Type | ICD-10 Classification | Tissue | |

Through browsing the database or after a search is submitted, a list of diseases appears, as the shown above. The list contains the Disease Name, the ICD-10 Classification, the disease type and the tissue(s), in which deposits are located. When the user presses the Show button they are redirected to the Entry page of a Disease.

#### Disease Entry

The Entry page contains information about the disease. CytoscapeJS is integrated to visualize the relationship between the disease and proteins found on amyloid deposits.

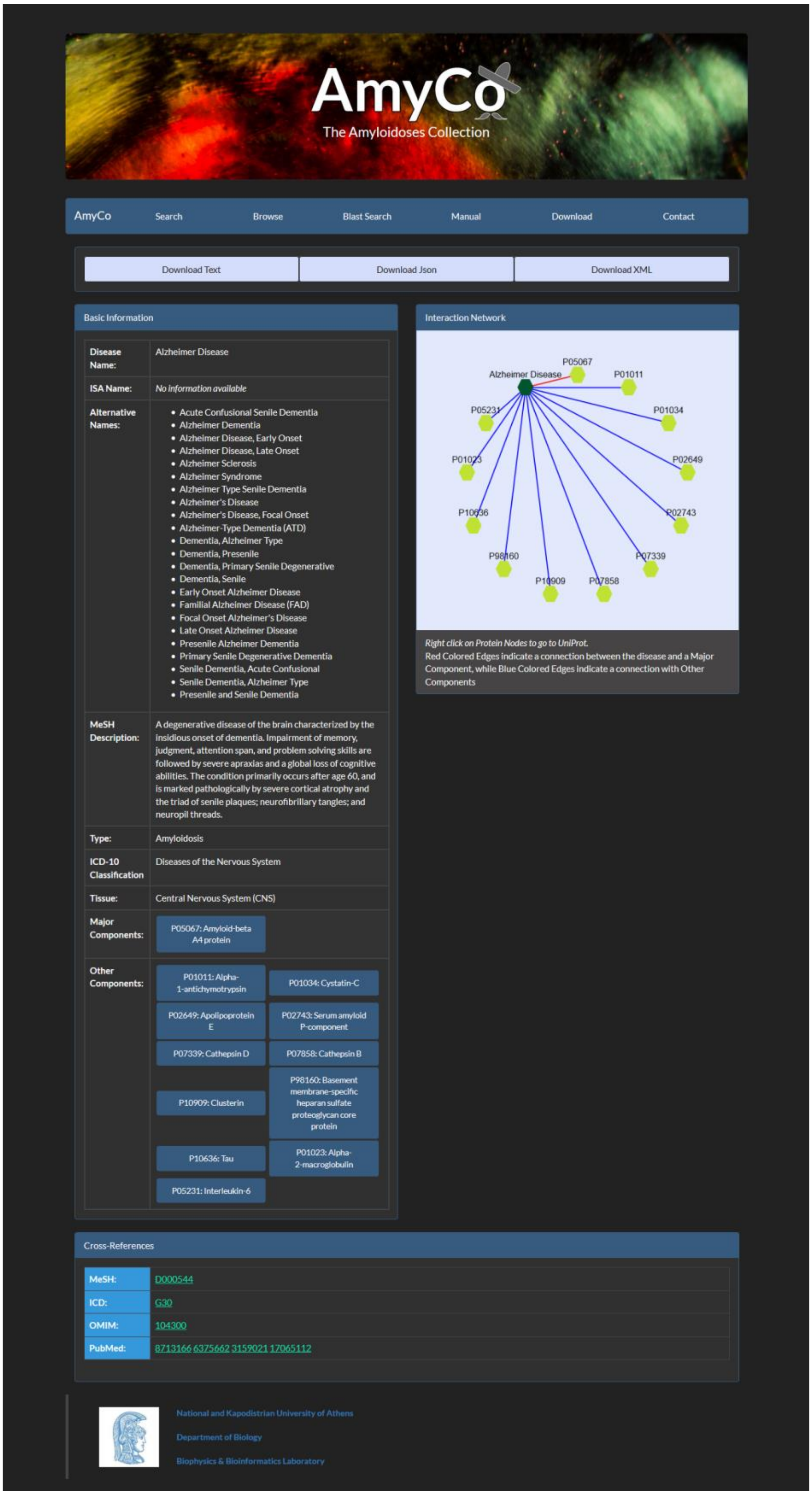

At the top of the page the user can find three buttons; *Download Text*, *Download Json* and *Download XML*. By pressing these buttons the user can download all page information in text, Json or XML format respectively.

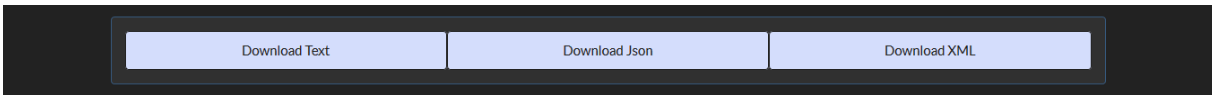


The basic disease information available is:

- ✓ Disease Name (MeSH name, when the disease is a MeSH entry)
- ✓ ISA Name (International Society of Amyloidosis name, when available)
- ✓ Alternative Names
- ✓ Disease Description (a short description from MeSH)
- ✓ DiseaseType (see here)
- ✓ Disease Association (from ICD-10, when available)
- ✓ Tissue(s) where amyloid deposits are located
- ✓ Major Components of Amyloid Deposits
- ✓ Other Components of Amyloid Deposits

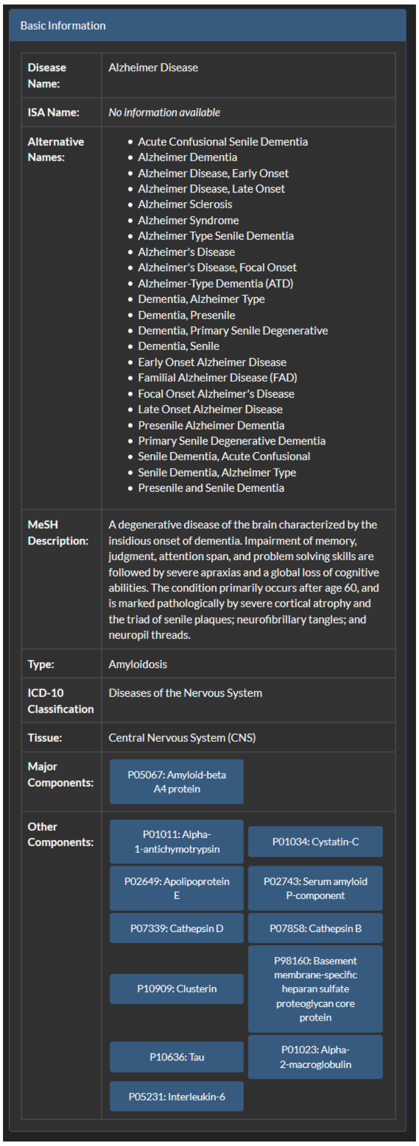

The panel at the bottom of the page contains references to other databases. The databases are:

- MeSH
- ICD
- OMIM
- PubMed

Cross-References

| | |
|---|---|
| MeSH: | D000544 |
| ICD: | G30 |
| OMIM: | 104300 |
| PubMed: | 20061647 |

When the user presses on the buttons of major or other components a new page opens with information about the proteins.

- Primary Name
- Gene Name
- Other Protein Names
- Protein Sequence
- Protein Length
- Uniprot AC
- Uniprot ID

For major components an external link to AmyPro is given when available. For other components the peer reviewed publication (*Association Source*) of disease is also provided.

Protein Information

| | |
|---|---|
| Primary Name: | Cystatin-C |
| Gene Name: | CST3 |
| Association Source: | 11202179 |
| Protein Names: | Cystatin-C (Cystatin-3) (Gamma-trace) (Neuroendocrine basic polypeptide) (Post-gamma-globulin) |
| Protein Length: | 146 |
| Protein Sequence: | MAGPLRAPLLLLAILAVALAVSPAAGSSPGKPPRLVGGPMDASVEEEGVRRALDFAVGEYNKASNDMYHSRALQVVRARKQIVAGVNYFLDVELGRTTCTKTQPNLDNCPFHDQPHLKRKAF CSFQIYAVPWQGTMTLSKSTCQDA |
| UniProt AC: | P01034 |

A CytoscapeJS viewer is integrated in the page for the visualization of bipartite graphs, showing the association between **each disease** with **major and other protein components**.

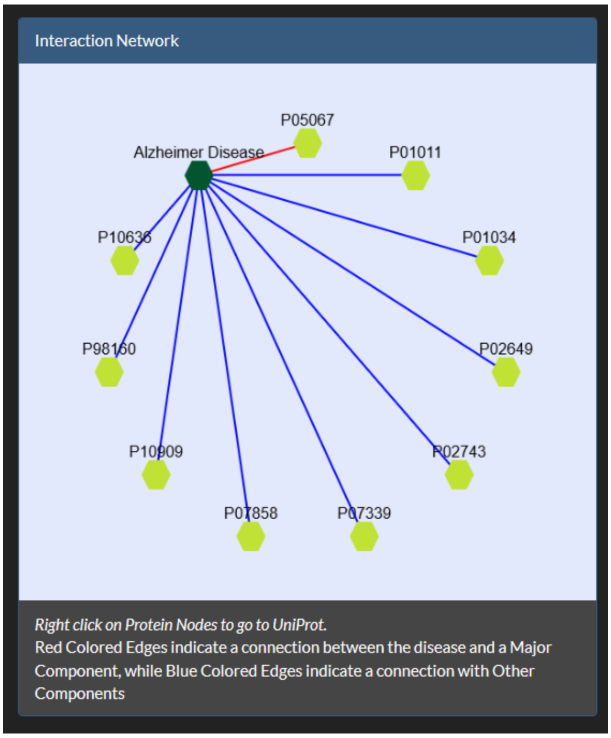


The diseases are colored green and the protein components yellow. Red Colored Edges indicate a connection between the disease and a Major Component, while Blue Colored Edges indicate a connection with Other Protein Components. Each protein node is also a hyperlink to UniProt.

If your browser prevents you from opening pop-up windows, please select *Allow pop-ups for this site*.

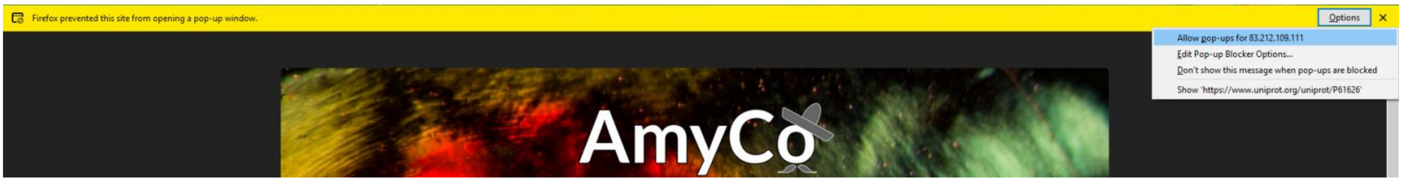


If during scrolling through the page you accidentally lose the network view, please reload the page to see it again.

#### BLAST Search

With the BLAST search tool, the user may submit a sequence and search the database for homologous protein molecules.

The input for the BLAST application is a sequence in the standard FASTA format.

```
>sp|P01034|27-146
SSPGKPPRLVGGPMDASVEEEGVRR
```

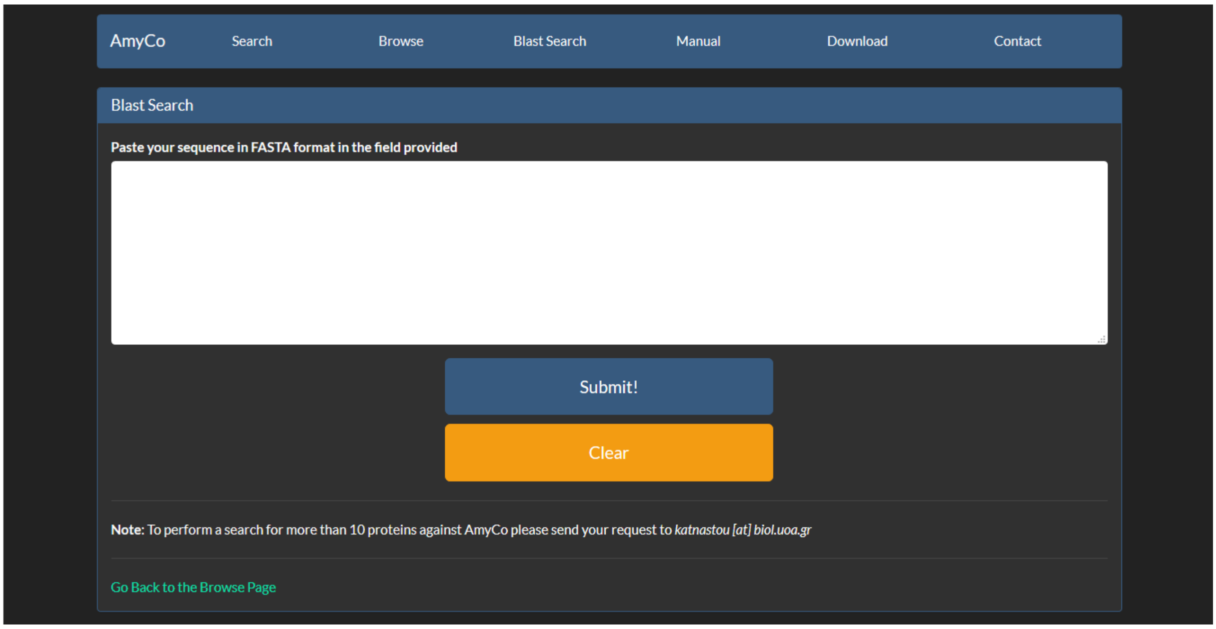


The result page of the BLAST search shows a list of the Blast hits with significant alignment on the query sequence. The list is in a table format including the target protein, the Length of the target sequence and the Query and finally, Target align range.

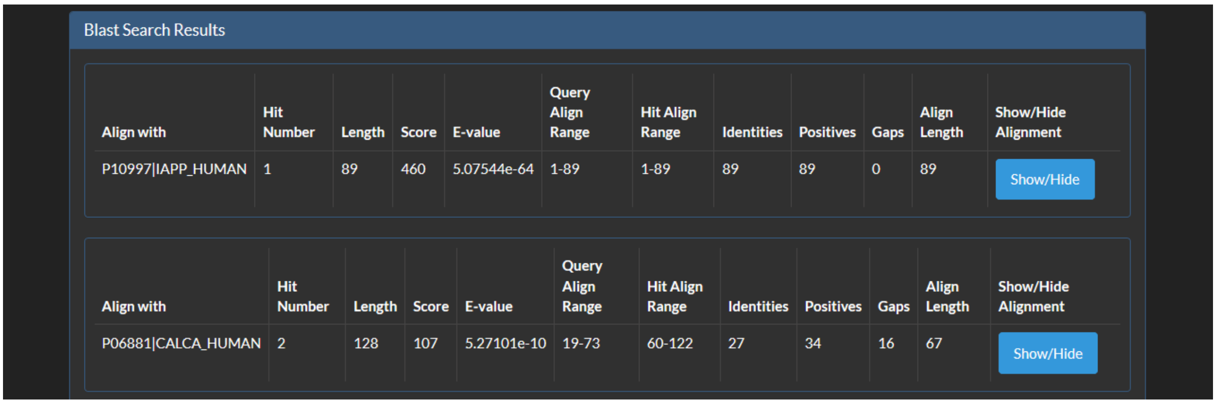


The BLAST results can be compared through the Score, the E-value, the Identities and Positives.

Furthermore, the user can have a more detailed view of each alignment through the Show button at the end of each line:

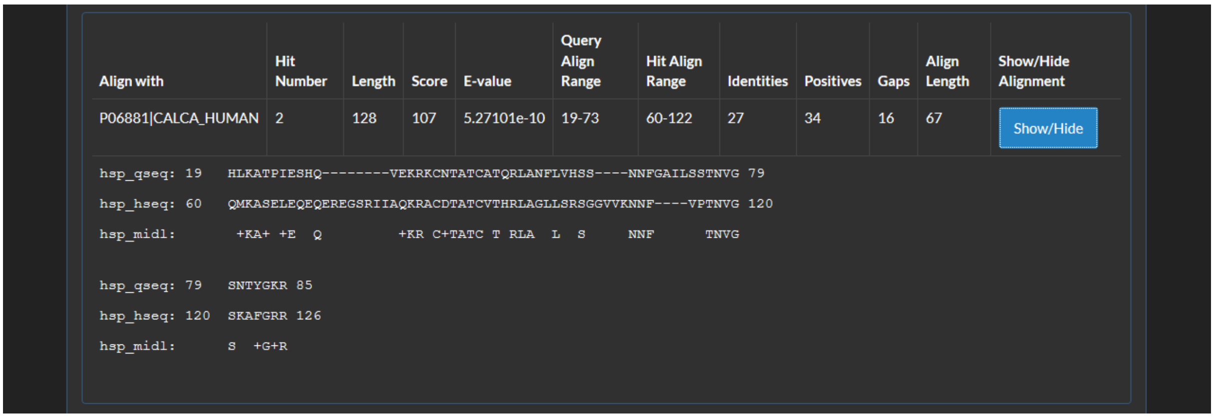


#### Download

User can download all database files in Text, JSON or XML format.

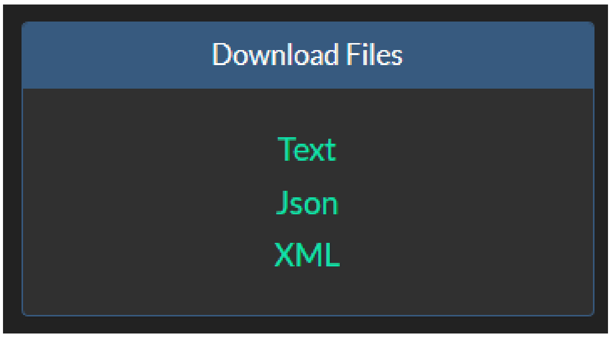


#### Contact

Users can contact us for more information at the emails specified at the contact page.

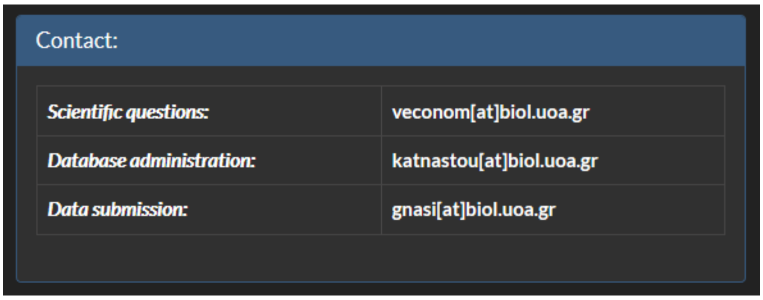

Users are encouraged to submit data by using the form below.  Data will be reviewed and later will be added to the database by the authors.

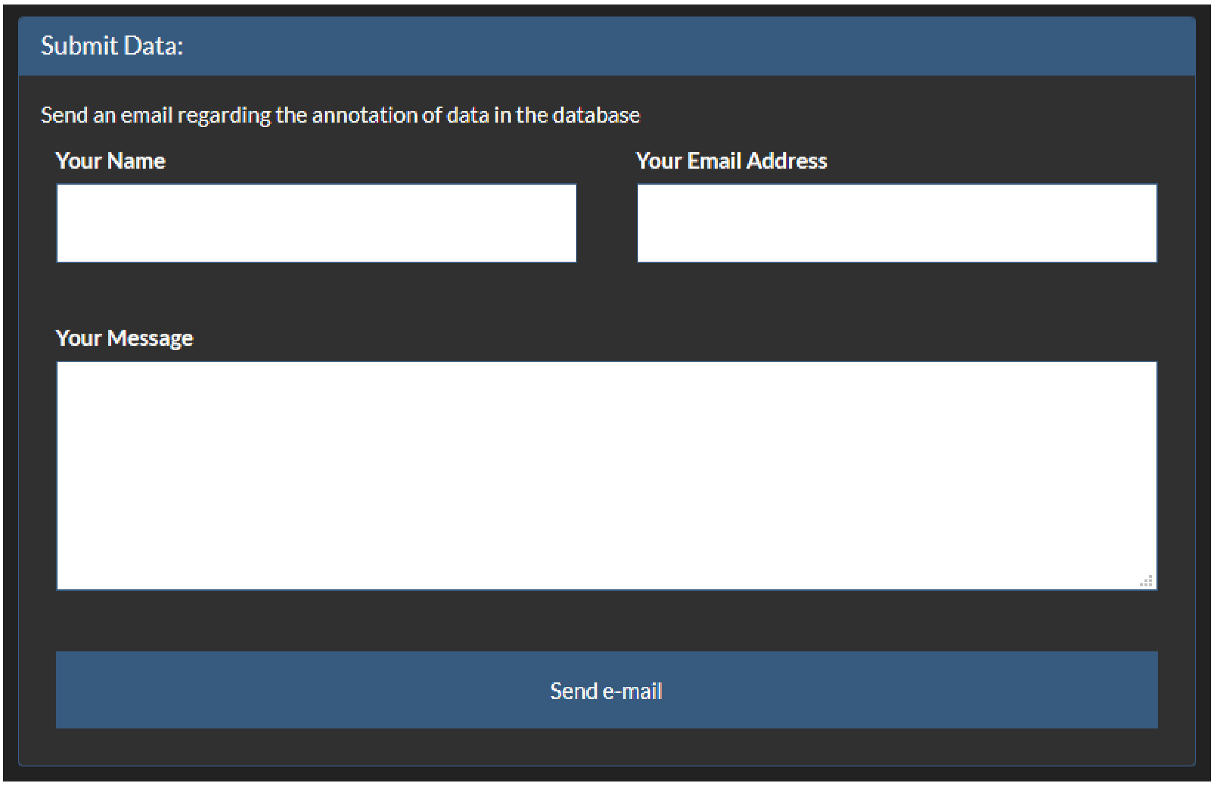
Submit Data:

Send an email regarding the annotation of data in the database

Your Name

Your Email Address

Your Message

Send e-mail

Related publications to the current work are also presented.

#### Database Technologies

*AmyCo* is based on modern technologies. User should have Javascript enabled on the web browser.
